# Endometrium-on-a-chip reveals the endometrial transcriptome, and protein content of secretome are altered by changes in circulating concentrations of insulin and glucose *in vitro*

**DOI:** 10.1101/2020.11.03.361774

**Authors:** Tiago H. C. De Bem, Haidee Tinning, Elton J. R. Vasconcelos, Dapeng Wang, Niamh Forde

## Abstract

The molecular interactions between the maternal environment and developing embryo that are key for early pregnancy success are known to be influenced by factors such as the metabolic status. We are, however, limited in our understanding of the mechanism by which these individual nutritional stressors alter endometrial function and the *in utero* environment for early pregnancy success. Here we report for the first time the use of endometrium-on-a-chip microfluidics approach to produce a multi-cellular endometrium *in vitro*, that is exposed to glucose and insulin concentrations associated with maternal metabolic stressors. Following isolation of endometrial cells (epithelial and stromal) from the uteri of non-pregnant cows in early-luteal phase (Day 4-7 approximately) epithelial cells were seeded into the upper chamber (4-6 10^4^ cells/mL) and stromal cells seeded in the lower chamber (1.5-2 10^4^ cells/mL). Three different concentration of glucose 1) 0.5 mM 2) 5.0 mM or 3) 50 mM or insulin 1) Vehicle, 2) 1 ng/mL or 3) 10 ng/mL were performed in the endometrial cells at a flow rate of 1µL/min for 72 hr to mimic the rate of secretion *in vivo*. Quantitative differences in the transcriptomic response of the cells and the secreted proteome of *in vitro*-derived uterine luminal fluid (ULF) were determined by RNA-sequencing and *TMT* respectively. Changes in maternal glucose altered 21 and 191 protein coding genes in epithelial and stromal cells respectively (p<0.05). While there was a dose-dependent quantitative change in protein secretome (1 and 23 proteins). Insulin resulted in limited transcriptional changes including insulin-like binding proteins that were cell specific (5, 12, and 20) but altered the quantitative secretion of 196 proteins including those involved in extracellular matrix-receptor interaction and proteoglycan signaling in cancer. Collectively, these highlight the potential mechanism by which changes to maternal glucose and insulin alter uterine function.

## INTRODUCTION

Successful early pregnancy in placental mammals requires bilateral interactions between the developing embryo and the maternal endometrium. While interaction with the maternal environment is not strictly required, i.e. we can produce embryos *in vitro* that give rise to healthy offspring, interactions with the maternal tract substantially enhances the quality of the embryo (1–3). This increased developmental competency is mediated in part via the transport and secretion of endometrial derived molecules (including proteins, amino acids, metabolites, lipids and RNA species encapsulated in extracellular vesicles (EVs)) that are taken up by the embryo to support development prior to establishment of the placenta (4, 5). The composition of uterine luminal fluid (ULF) and the molecular interactions between the mother and developing embryo are known to be influenced by maternal factors such as the metabolic status of the mother as well as the quality of the embryo present (Reviewed by (6, 7)). Exposure to adverse conditions e.g. nutritional insults, at specific developmental time points can alter an individual’s susceptibility to disease in later life (7). Despite advances in our understanding of the ULF composition (8, 9), efforts to supplement culture media with these components have not substantially improved development indicating there are likely still unknown components of ULF yet to be discovered.

The uterine epithelium, at least for a few days, is potentially the most critical maternal tissues in the establishment of a healthy pregnancy (10). Thus, exposure of the endometrium to stressors can alter the developmental or epigenetic programming of the foetus. In the dairy cow, there is a period of time post-partum, when they are under metabolic stress and cannot take in sufficient food to off-set the energy demands of peak milk production. This induces a maternal metabolic environment that has high Non-Esterified Fatty Acids, Beta Hydroxy Butaric Acid, and low insulin, IGF-I, and glucose (11, 12). This occurs co-ordinately with establishing early pregnancy, and there is a growing body of evidence to suggest that this metabolic stress in maternal circulation compromises the ability of the reproductive tract of the lactating dairy cow to support early development (11, 12). Data indicates this metabolic stress alters global gene expression in the embryo/conceptus, the oviduct (9, 13), and endometrium (13–15). Conceptuses derived from early in lactation are less developmentally competent (metabolic stress) compared to late stages of lactation (16). Even if a high-quality embryo is produced exposure to the uterine environment of lactating cows can alter developmental potential (16, 17).

We are, however, limited in our understanding of the mechanism by which these individual components alter endometrial function and the *in utero* environment. While traditional static cell culture models have been used to address some of these conceptus-maternal interactions they do have limitations they do not mimic the continual flow of these metabolic components to which the endometrium is exposed. Nor do they allow for assessment of how these metabolic extremes alter the interactions between the heterogenous cells types of the endometrium and the ULF that is produced as a consequence. Advances in microfluidics and organ-on-a-chip technologies in reproductive systems include accessing embryo culture systems (18), cervical (19), ovarian (20), endometrial (21), and placental function (22, 23) in humans and mice. There have been reports of using these systems to mimic the bovine oviduct environment (24) and there has been the breakthrough publication of modelling of the menstrual cycle *in vitro* using a combination of human and murine components (25). However, no one has harnessed these systems to investigate how maternal nutritional stressors alter the uterine environment to which the embryo is exposed either at the level of the transcriptomic or proteomic secretome.

Here we report for the first time the use of microfluidics to produce a multi-cellular endometrium *in vitro*, that is exposed to glucose and insulin concentrations associated with maternal metabolic stressors. We used RNA sequencing to determine how these metabolites alter the cell-specific transcriptional responses in the endometrium to these nutritional stressors. We further demonstrate how these changes alter the proteomic content of *in vitro*-derived ULF secreted from the endometrial epithelium. Collectively, these highlight the potential mechanism by which changes to maternal glucose and insulin alter uterine function. We propose that these are candidate molecules that can modify the developmental potential of embryos.

## MATERIALS AND METHODS

Unless otherwise stated, all chemical and reagents were purchased from Sigma-Aldrich Chemical Co. (St. Louis, MO). The *in vitro* experimental procedures were conducted in humidified incubators maintained at 38.5°C with 5% CO_2_ in air.

### Primary Endometrial Cell Isolation and Culture

Endometrial cell isolation was carried out as previously described (26). Briefly, uteri from non-pregnant cows (*Bos taurus*), early in the luteal phase (Day 4-7 approximately) were selected on the basis of *corpus luteum* morphology as described (27). Endometrial tissue was dissected from the underlying myometrium and incubated in 25 mL digest solution, containing bovine serum albumin (1 mg/mL, BSA), trypsin EDTA (2.5 BAEE units/mL), collagenase (0.5 mg/mL) and DNase I (0.1 mg/mL) in Hanks Buffered Saline Solution (HBSS) in a shaking hot bath for 1 hr at 37 °C. The digestive solution was filtered through a 100μm mesh cell strainer over a 4μm cell strainer, washed twice with HBSS (containing 10% FBS), and centrifuged at 700 g for 7 min. The resulting cell pellet containing stromal cells was re-suspended in RPMI 1640 culture media containing 10% FBS, streptomycin (50 μg/mL), and penicillin (50 IU/mL) amphotericin B (2.5 μg/mL). The 40μm strainer was inverted and flushed with culture media to recover epithelial cells. The cell populations were seeded at 1×10^5^ cells/mL into 75 cm^2^ culture flasks (Greiner BioOne, Gloucestershire, UK). After 5-7 days of culture cell populations were further purified using their differential plating times.

### Cell Seeding into the Microfluidic Device

All the cells were seeded into the devices in RPMI 1640 media as described above (Figure 1). Stromal cells were seeded in the lower chamber of the devices (10µm-slide membrane, IBIDI), using a 1 mL syringe, at concentration of 1.5-2×10^4^ cells/mL in a final volume of 300 µL. All devices were inverted for 2 hr to allow stromal cells to adhere to the underside of the porous membrane. Devices were placed in the normal orientation and epithelial cells seeded at 4-6×10^4^ cells/mL in a final volume of 55 µL into the upper chamber. Cells were left to become 60% confluent over two days with one media change (48 hr) before beginning the flow perfusion. For the Glucose experiment, on the day of experimentation media was changed and 5 mL of media without glucose was loaded into 5mL syringes at the inlet (n = 2 technical replicates) with either 1) 0.5 mM, 2) 5.0 mM, or 3) 50.0 mM of glucose. For the insulin experiment, on the day of experimentation media was also changed and 5 mL of media with physiologic concentration of glucose (5.0 mM) was loaded into 5 mL syringes at the inlet (n = 2 technical replicates) with either 1) Vehicle control (acetic acid in PBS pH3.0), 2) 1.0 ng/mL or 3) 10.0 ng/mL of insulin. The pump was set to flow media through the device at a flow rate of 1µL/min to mimic the rate of secretion *in vivo* (28) for 72 hr. Culture media from the upper chamber (*in vitro*-derived ULF) was recovered with a pipette and snap-frozen in liquid nitrogen and the samples were stored at −80 °C. Epithelial and stromal cells were separately removed from the device via 0.5% trypsin, centrifuged at 2000 g for 10 min, snap-frozen in liquid nitrogen, and stored at −80 °C prior to processing.

**Figure 1.**
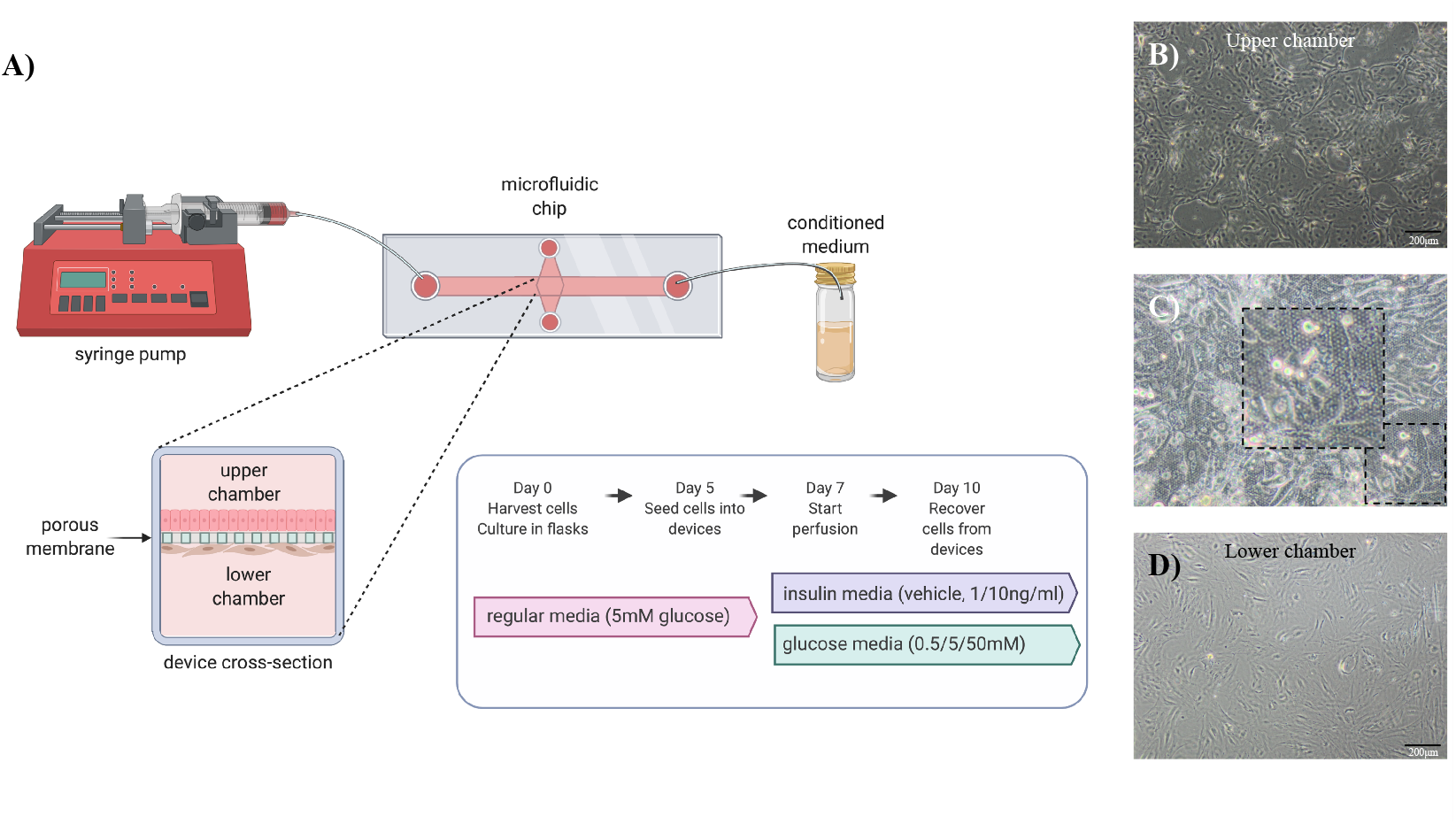
**A)** Schematic diagram of endometrium on-a-chip microfluidic device and experimental design used to mimic the physiological extremes of glucose and insulin. Representative images of **B)** epithelial cells seeded into the upper chamber, **(C)** cells attached on and under the porous membrane, and **D)** stromal cells seeded into the lower chamber. The rate of flow for both experiments was performed at 1uL/min per 72h.

### RNA extraction and sequencing

Total RNA was extracted from epithelial and stromal cells using the Mini RNeasy kit (Qiagen, Crawley, UK) following the manufacturer’s recommendations. Cell samples were homogenized in 700 µL of Qiazol via vortexing for 1 min at RT. On-column DNase digestion was performed (15 min at room temperature (RT)) and RNA was eluted in 14 µL of RNase/DNase free water from the spin column membrane following centrifugation for 2 min at full speed and this step was performed twice.

RNA sequencing was performed as previously described (26) with minor modifications. Briefly, RNA quality and quantity were confirmed using the Agilent Bioanalyzer system, and all samples had an RNA integrity number of >7.9. Stranded RNA sequencing libraries were constructed using the Encore Complete RNA-Seq library system of NuGEN. All libraries were sequenced on the Illumina HiSeq 1500 generating NextSeq 75 bp single-endend reads per library. The raw FASTQ files were inspected using FastQC (https://www.bioinformatics.babraham.ac.uk/projects/fastqc/), the adapter sequences were trimmed using Cutadapt and additional quality control steps taken by fastq_quality_filter program as part of FASTX-Toolkit (http://hannonlab.cshl.edu/fastx_toolkit/). The mapping process was performed using align function in Rsubread package by aligning the clean fastq files against the cow reference genome retrieved from Ensembl release 96 (*Bos taurus*) and only uniquely-mapped alignments were recorded. The resulting BAM files were sorted and indexed by SAMtools. The reads were summarized at the gene level by means of featureCounts. DESeq2 in a paired sample design was used to identify differentially expressed protein-coding genes based on the cut-offs of a p value < 0.05 and a log_2_ Fold Change > 0.1 or < −0.1. Variance stabilizing transformations were applied to the genes that have at least 10 reads in total for all samples. Heatmaps were created using the transformed read counts based on a pool of differentially expressed protein-coding genes for each experiment. PCA analysis was carried out for the protein-coding genes that have RPKM >= 1 in at least one sample and two normalisation approaches such as log2(RPKM+1) and quantile normalization were conducted prior to PCA analysis.

### Quantitative proteomic analysis of in vitro-derived uterine luminal fluid recovered from the upper chamber

Media from the upper chamber (*in vitro*-derived ULF) from glucose 0.5 mM (n = 3), 5 mM (n = 3) and 50 mM (n = 3) and insulin Vehicle (n = 3), 1 ng/mL (n = 3) and 10 ng/mL (n = 3) treatment were subjected to albumin depletion according to the manufacturer’s instructions (Thermo Fisher Scientific, Loughborough, UK). Individual samples were digested with trypsin (2.5µg trypsin; 37 °C, overnight), labelled with Tandem Mass Tag (TMT) ten plex reagents according to the manufacturer’s protocol (Thermo Fisher Scientific) and pooled. The pooled sample was evaporated to dryness, resuspended in 5% formic acid, and then desalted using a SepPak cartridge according to the manufacturer’s instructions (Waters, Milford, Massachusetts, USA). Eluate from the SepPak cartridge was again evaporated to dryness and resuspended in buffer A (20 mM ammonium hydroxide, pH 10) prior to fractionation by high pH reversed-phase chromatography using an Ultimate 3000 liquid chromatography system (Thermo Scientific). In brief, the sample was loaded onto an XBridge BEH C18 Column (130Å, 3.5 µm, 2.1 mm X 150 mm, Waters, UK) in buffer A and peptides eluted with an increasing gradient of buffer B (20 mM Ammonium Hydroxide in acetonitrile, pH 10) from 0-95% over 60 min. The resulting fractions were evaporated to dryness and resuspended in 1% formic acid prior to analysis by nano-LC MSMS using an Orbitrap Fusion Lumos mass spectrometer (Thermo Fisher Scientific).

### Nano-LC Mass Spectrometry

High pH reversed-phase (RP) fractions were further fractionated using an Ultimate 3000 nano-LC system in line with an Orbitrap Fusion Lumos mass spectrometer (Thermo Scientific). Peptides in 1% (vol/vol) formic acid were injected onto an Acclaim PepMap C18 nano-trap column (Thermo Scientific). After washing with 0.5% (vol/vol) acetonitrile 0.1% (vol/vol) formic acid peptides were resolved on a 250 mm × 75 μm Acclaim PepMap C18 reverse phase analytical column (Thermo Scientific) over a 150 min organic gradient, using 7 gradient segments (1-6% solvent B over 1 min, 6-15% B over 58 min, 15-32% B over 58 min, 32-40% B over 5 min, 40-90% B over 1min, held at 90% B for 6 min and then reduced to 1% B over 1 min) with a flow rate of 300 nl min−1. Solvent A was 0.1% formic acid and Solvent B was aqueous 80% acetonitrile in 0.1% formic acid. Peptides were ionized by nano-electrospray ionization at 2.0kV using a stainless-steel emitter with an internal diameter of 30 μm (Thermo Scientific) and a capillary temperature of 275 °C. All spectra were acquired using an Orbitrap Fusion Lumos mass spectrometer controlled by Xcalibur 4.1 software (Thermo Scientific) and operated in data-dependent acquisition mode using an SPS-MS3 workflow. Fourier transform mass analysers 1 (FTMS1) spectra were collected at a resolution of 120 000, with an automatic gain control (AGC) target of 200 000 and a max injection time of 50ms. Precursors were filtered with an intensity threshold of 5000, according to charge state (to include charge states 2-7) and with monoisotopic peak determination set to Peptide. Previously interrogated precursors were excluded using a dynamic window (60s +/-10ppm). The MS2 precursors were isolated with a quadrupole isolation window of 0.7m/z. ITMS2 spectra were collected with an AGC target of 10 000, max injection time of 70ms and CID collision energy of 35%. For Fourier transform mass analysers 3 (FTMS3) analysis, the Orbitrap was operated at 50 000 resolution with an AGC target of 50 000 and a max injection time of 105ms. Precursors were fragmented by high energy collision dissociation (HCD) at a normalised collision energy of 60% to ensure maximal TMT reporter ion yield. Synchronous Precursor Selection (SPS) was enabled to include up to 5 MS2 fragment ions in the FTMS3 scan.

The raw data files were processed and quantified using Proteome Discoverer software v2.1 (Thermo Scientific) and searched against the UniProt *Bos taurus* database (downloaded June 2019: 46309 entries) using the SEQUEST algorithm. Peptide precursor mass tolerance was set at 10ppm, and MS/MS tolerance was set at 0.6Da. Search criteria included oxidation of methionine (+15.9949) as a variable modification and carbamidomethylation of cysteine (+57.0214) and the addition of the TMT mass tag (+229.163) to peptide N-termini and lysine as fixed modifications. Searches were performed with full tryptic digestion and a maximum of 2 missed cleavages were allowed. The reverse database search option was enabled, and all data was filtered to satisfy false discovery rate (FDR) of 5%.

### Regulatory networks and functional annotations of biological processes analysis were performed by bioinformatics using DAVID

Gene enrichment analysis was performed using DAVID (Database for annotation, visualization, and integrated discovery; Bioinformatics Resources 6.7) (29) to predict the regulatory networks (signalling pathways) and specific functional annotations from gene ontology (GO) terms related to the proteins and DEGs, respectively. The accession number from each protein (Uniprot) or gene (Ensembl Gene ID) were used. All the signalling pathways GO terms identified from biological process, cellular component, and molecular function were considered enriched a P<0.05 threshold. To perform and visualization of the networks of the signalling pathways was used the software platform Cytoscape version 3.7.2 (30). The signalling pathways were presented by the Fold Enrichment each networking and the GO terms were arranged by the Enrichment Score (-log_10_^[Pvalue]^), respectively.

## RESULTS

### Exposure of endometrial epithelial and stromal cells in vitro to high concentrations of glucose alters the transcriptional profile in a cell specific manner

Principal Component Analysis (PCA: Figure 2) plot showed distinct clustering of epithelial and stromal cells but there was no clear clustering between the treatments of glucose (5mM vs 50mM). Flow of high concentrations of glucose (50mM) for 3 days altered the expression of 21 and 191 protein coding genes compared to control (5.0 mM glucose) in the epithelial and stromal cells, respectively (Supplementary Table 1). Functional annotation analysis determined n = 8 and n = 14 overrepresented biological processes, n = 6 and n = 5 overrepresented cellular components and n = 4 and n = 6 overrepresented molecular functions, for epithelial and stromal cells exposed to 50 mM glucose, respectively (Figure 3; Supplementary Table 2).

**Figure 2.**
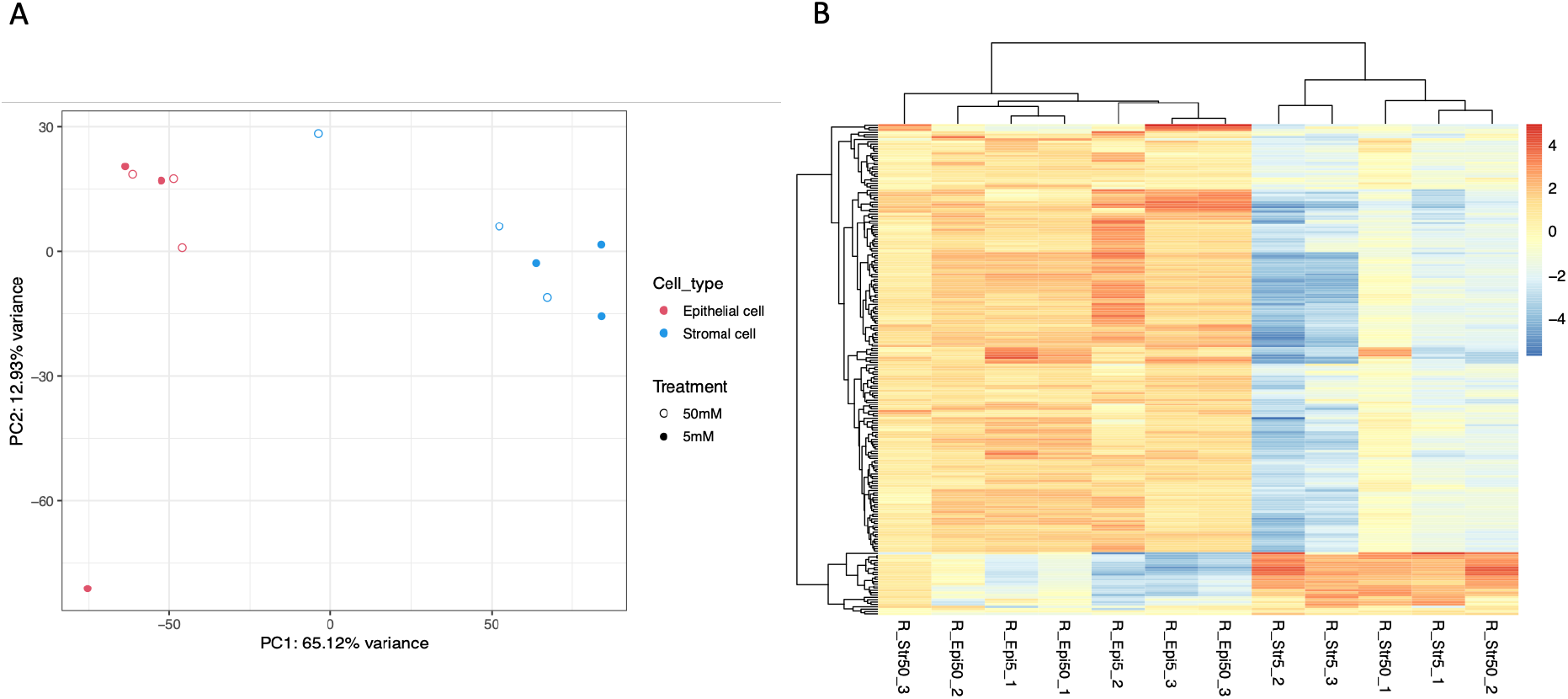
**A)** Principal Component Analysis (PCA) plot of overall transcriptional profile determined via RNA sequencing of bovine endometrial epithelial and stromal cells exposed to either 5 mM or 50 mM of glucose for 72 hr under flow (n=3 biological replicates). Epithelial (left hand side) and stromal cells (right hand side) clustered into two distinct populations. **B)** Heatmap showing the transcript expression for individual samples with lower levels (blue) and those with higher expression shown in red. Samples from stromal cells (n = 9) and samples from epithelial cells (n = 9) are arranged from left to the right.

**Figure 3.**
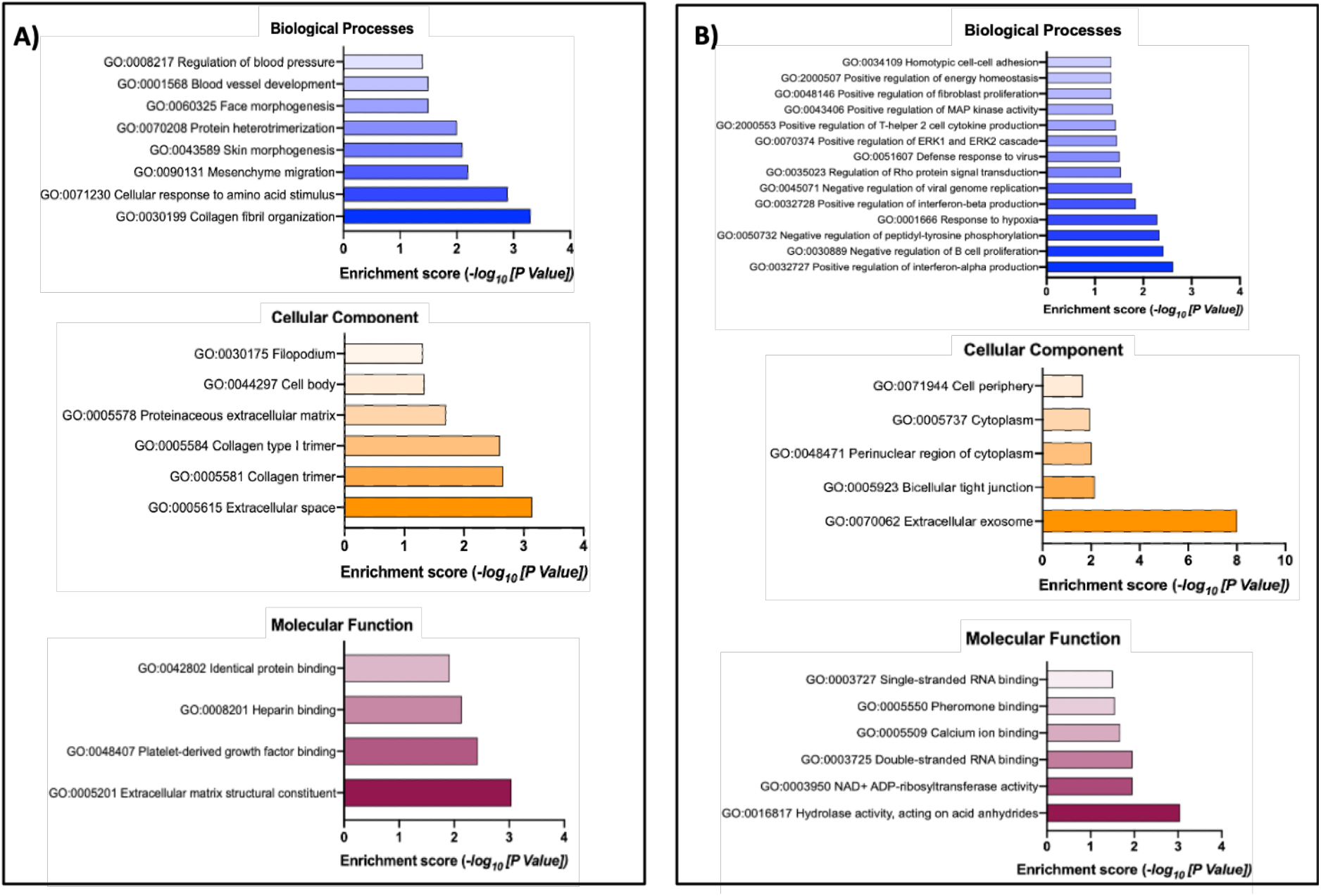
Gene Ontology overrepresented Biological Process, Cellular Component, and Molecular Function from DEGs in **A)** epithelial (n=21) and, **B)** stromal cells (n=191) following exposure to different concentrations of glucose *in vitro*. The transcript accession numbers (Ensembl Transcript ID) were inputted into DAVID - Functional Annotation Tool (DAVID Bioinformatics Resources 6.7, NIAID/NIH - https://david.ncifcrf.gov/summary.jsp) and those that were significantly overrepresented in the list of DEGs are presented above for biological processes (blue bars), cellular component (orange bars) and molecular function (plum bars) with associated enrichment score.

### Exposure to high concentrations of glucose alters the proteomic content of in vitro-derived ULF

PCA plot (Figure 4A) revealed that exposure of endometrium on-a-chip to physiological extremes of glucose (0.5mM and 50mM) altered the overall composition of proteins in the *in vitro*-derived ULF. The highest concentration of glucose (50mM) changed the abundance of 23 proteins compared to controls (5mM), the majority of which were increased (p < 0.05: Figure 4B). When the lower concentration was compared with the control (0.5mM vs 5mM), only one protein was altered. Finally, when the physiologic extremes (0.5mM vs 50mM) were compared, eight protein were found to be differentially abundant in *in vitro*-derived ULF. Functional annotation analysis revealed the proteins up-regulated following glucose treatment were involved in Platelet and Lysosome pathways (Supplementary Table 3). There were no over-represented pathways associated with the proteins that were decreased in abundance following glucose exposure (P>0.05).

**Figure 4.**
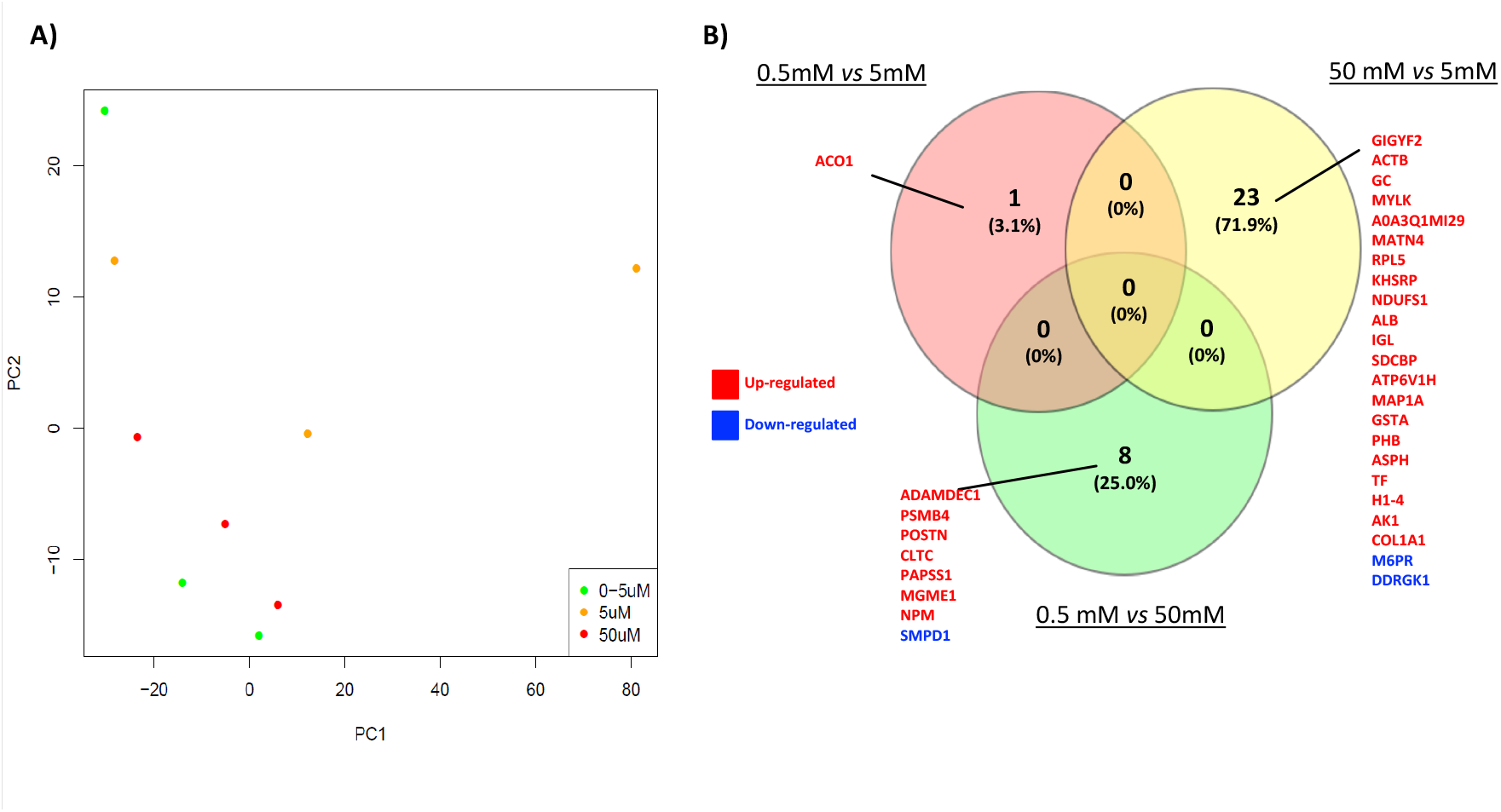
**A)** Principal component analysis (PCA) plot shows the distribution of *in vitro*-derived ULF following Tandem Mass Tag (TMT)-mass spectrometry analysis of the proteomic content following exposure to: 0.5mM (green circle), 5.0mM (yellow circle), or 50mM (red circle) concentrations of glucose for 72 hr (n=3 biological replicates). **B)** Venn diagram analysis of overlap and unique proteins that are differentially abundance following exposure to different concentrations of glucose. Fold change differences in protein abundances between groups was determined using paired t-tests and were considered significant when P<0.05.

### Exposure of the endometrium to different concentrations of Insulin alters the transcriptome in a cell specific manner

The PCA plot showed distinct clustering of epithelial cells on the left-hand side and stromal cells on the right with limited treatment effect (Figure 5A). Exposure of cells to 1 ng/mL of insulin for 72 hr changed expression of four transcripts (non-specific serine/threonine protein kinase (*ARAF*), Ubiquitin-40S ribosomal protein S27a (*RPS27A*), NADH-ubiquinone oxidoreductase chain 4L (*M-ND4L*), and NADH-ubiquinone oxidoreductase chain 6 (*MTND6*)) in epithelial cells and one unknown transcript (ENSBTAG00000052092) in stromal cells compared to vehicle control. Exposure to the higher concentrations of insulin (10 ng/mL) altered 10 transcripts in epithelial cells (NADH-ubiquinone oxidoreductase chain 2 (*MT-ND2*)), (NADH-ubiquinone oxidoreductase chain 3 (*MT-ND3*), (NADH-ubiquinone oxidoreductase chain 5 (*ND5*)), (CCAAT/enhancer-binding protein beta (*CEBPB*)), (Ferritin heavy chain 1(*FTH1*)), (ATP synthase protein 8 (*MT-ATP8*)), (Nuclear receptor subfamily 4 group A member 2 (*NR4A2*)), (Fibronectin (*FN1)*), (Collagenase 3 (*MMP13)*) and (Cytochrome b (*MT-CYB*)) and two transcripts in stromal cells: Insulin-like growth factor-binding protein 6 (*IGFBP6*), and receptor for retinol uptake (*STRA6*).

**Figure 5.**
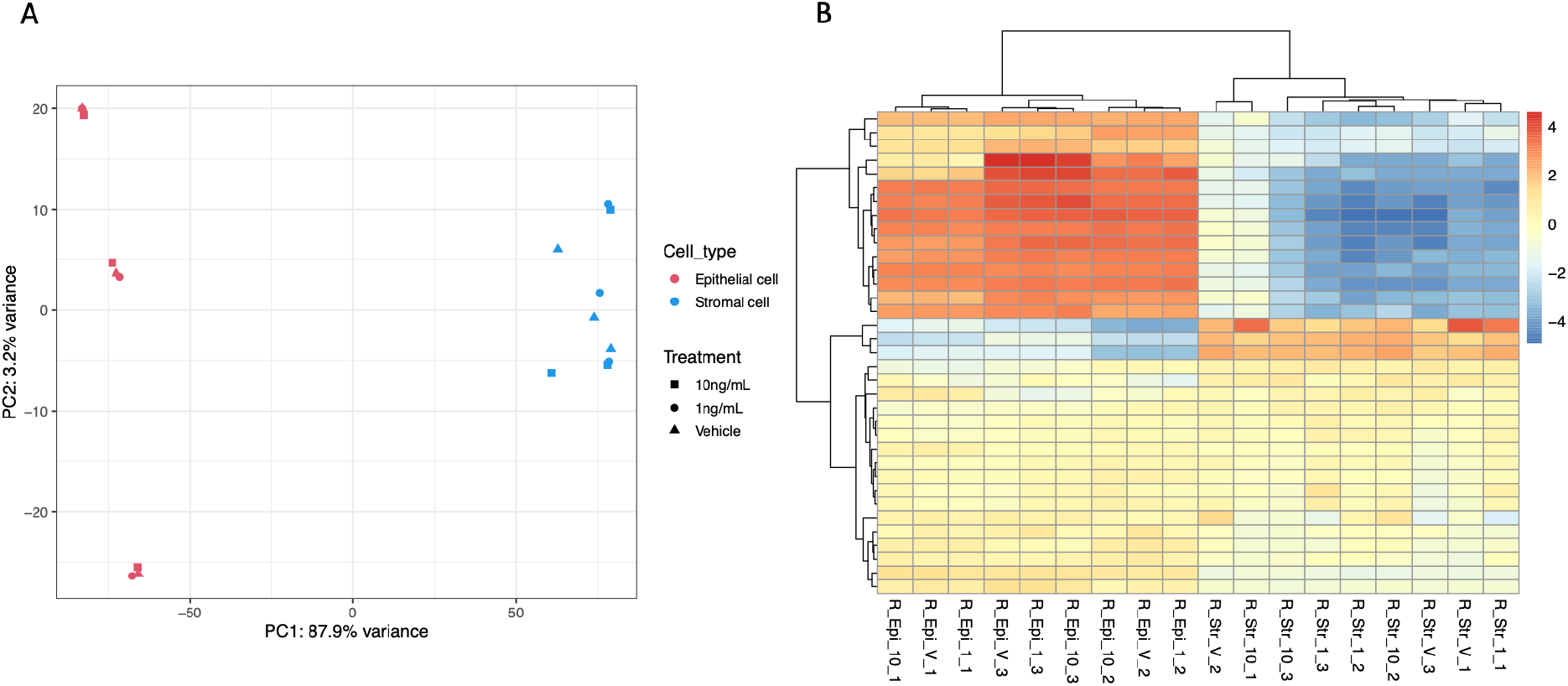
**A)** Principal Component Analysis (PCA) plot of overall transcriptional profile determined via RNA sequencing of bovine endometrial epithelial and stromal cells exposed to either vehicle control, 1 ng/mL, or 10 ng/mL of Insulin for 72 hr under flow (n=3 biological replicates). Epithelial (left hand side) and stromal cells (right hand side) clustered into two distinct populations. **B)** Heatmap showing the transcript expression for individual samples with lower levels (blue) and those with higher expression shown in red. Samples from stromal cells (n = 9) and samples from epithelial cells (n = 9) are arranged from left to the right.

In addition, when the physiological extremes of insulin (1ng *vs* 10ng/mL) were compared, one transcript was differentially expressed in epithelial cells (Insulin-like growth factor-binding protein 3 (*IGFBP3*)) while nineteen transcripts were altered in the stromal cells (Insulin-like growth factor-binding protein 5 (*IGFBP5*)), tensin 4 (*TNS4*), ETS domain-containing transcription factor (*EHF*), cytochrome P450, subfamily IIIA, polypeptide 4 (*CYP3A4*), sphingolipid delta(4)-desaturase (*DES1*), CCAAT/enhancer-binding protein beta, C/EBP beta (*CEBPB*), creatine kinase U-type, mitochondrial (*CKMT1A*), claudin-7 *(CLDN7*), mucin 16 (*MUC16*), glycine amidinotransferase, mitochondrial (*GATM*), claudin 6 (*CLDN6*), receptor protein-tyrosine kinase (*ERBB3*), ATP binding cassette subfamily A member 1 (*ABCA1*), epithelial membrane protein 1 (*EMP1*), solute carrier family 2, facilitated glucose transporter member 3 (*SLC2A3*), adseverin (*SCIN*), aldehyde dehydrogenase 1 family member A3 (*ALDH1A3*), vascular endothelial growth factor A (*VEGFA*), and NADH-ubiquinone oxidoreductase chain 6 (*MT-ND6*)). Functional annotation analysis found no overrepresented terms associated with the lower concentration of Insulin compared to control. However, DEGs associated with the higher concentration of insulin were overrepresetned in three cellular components (Respiratory chain, Mitochondrial respiratory chain complex I and Mitochondrial inner membrane) and one molecular function (NADH dehydrogenase (ubiquinone) activity).

### Altering insulin concentrations modifies the secretome of in vitro-derived ULF

PCA plot did not show distinct clustering in the overall proteomic profile of in vitro-derived ULF (Figure 6A). However, exposure of our endometrium on-a-chip to physiological extremes of insulin (1 ng/mL and 10 ng/mL) changed the abundance of 195 proteins (Figure 6B). The majority of these proteins (n = 67) were altered (P<0.05) in cells treated with the 1ng/mL of Insulin compared to vehicle control (Figure 6). The higher concentration of insulin altered 57 protein in total (P<0.05) while comparison of the two physiological extremes of insulin (1 ng/mL vs 10 ng/mL) revealed 51 differentially abundant proteins (P<0.05) proteins in *in vitro*-derived ULF. Venn diagram analysis showed that 17 proteins were altered in more than one group (Figure 6B). Proteins up-regulated following treatment with 1 ng/mL of Insulin were overrepresented in pathways associated with biosynthesis of amino acids (n = 2), carbon metabolism (n = 3), biosynthesis of antibiotics (n = 4), and metabolic pathways (n = 5). The down-regulated proteins were associated with complement and coagulation cascades (n = 3), protein processing in endoplasmic reticulum (n = 5), and amoebiasis (n = 3). Down-regulated proteins were overrepresented when cells were treated with 10 ng/mL of Insulin were related with protein digestion and absorption (n = 3), ECM-receptor interaction (n = 3) and proteoglycans in cancer (n = 4). All the pathways related with up or down-regulated proteins are presented in Figure 7.

**Figure 6.**
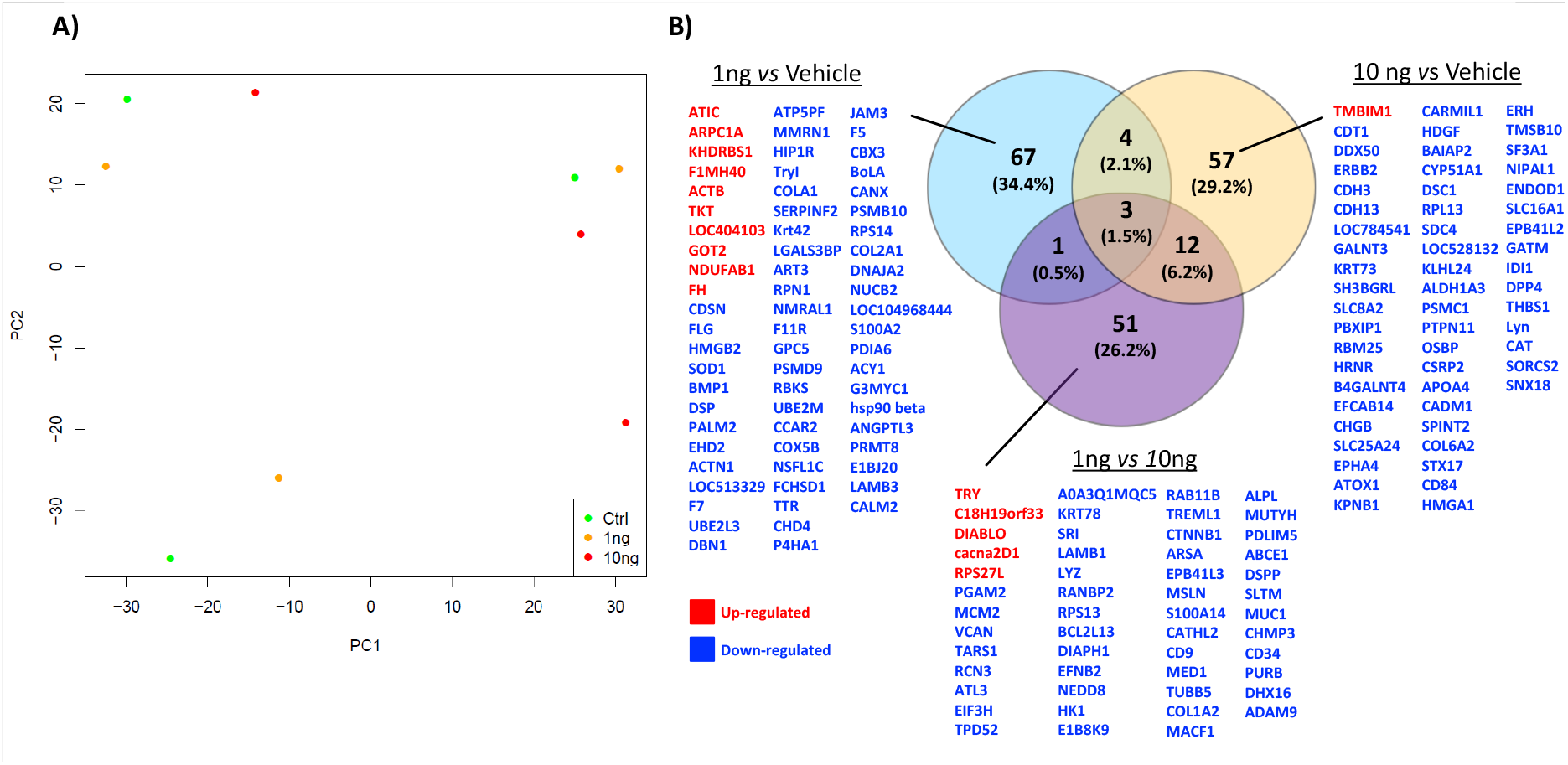
**A)** Principal component analysis (PCA) plot shows the distribution of *in vitro*-derived ULF following Tandem Mass Tag (TMT)-mass spectrometry analysis of the proteomic content following exposure to: vehicle control (green circle), 1 ng/mL Insulin (yellow circle), or 10 ng/mL Insulin (red circle) for 72 hr (n=3 biological replicates). **B)** Venn diagram analysis of overlap and unique proteins that are differentially abundant following exposure to different concentrations of insulin. Fold change differences in protein abundances between groups was determined using paired t-tests and were considered significant when P<0.05.

**Figure 7.**
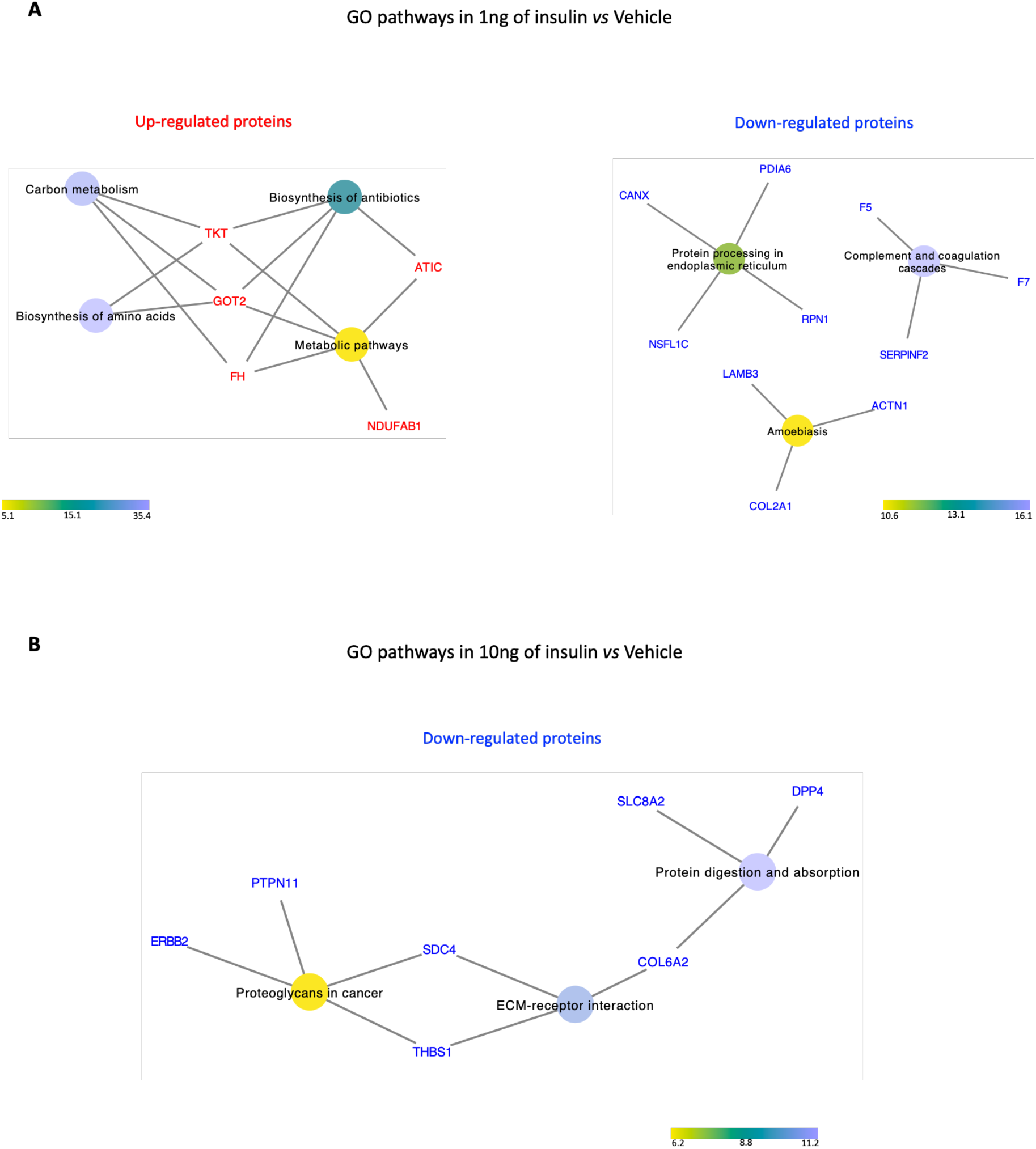
Network interaction between the differentially abundant proteins and their signaling pathways (P<0.05) from *in vitro*-derived ULF from endometrium on-a-chip exposed to insulin for 72 hr. **A)** GO pathways associated with proteins altered in abundance following treatment with 1.0 ng/mL of insulin compared to vehicle control. **B)** GO pathways associated with proteins altered in abundance following treatment with 10.0 ng/mL of insulin was compared to vehicle control. The color circle represents an individual signaling pathway and the proteins around it are proteins identified as altered in our treatments associated with that signaling pathway (s) what are either up-regulated (red) or down-regulated (blue) following insulin treatment. The color scale represents the Fold Enrichment that reflects the involvement of the proteins in each signaling pathway.

## DISCUSSION

This study aimed to generate a new *in vitro* model to study the interactions between the maternal metabolic environment and endometrial function. Using for the first time a microfluidics approach to mimic the bovine endometrium *in vitro*, we have identified the transcriptomic pathways that are changed by different physiologically relevant concentrations of glucose and insulin as well as the consequences for the secreted proteomic composition of the uterine luminal fluid produced *in vitro*. These supply us with evidence for the potential mechanism of action of two key metabolic stressors (glucose and insulin) independently on uterine function and begin to allow us to understand how maternal nutritional stressors contribute to uterine dysfunction and ultimately early pregnancy loss.

Concentrations of glucose and insulin are decreased in circulation in dairy cows undergoing negative energy balance (31, 32). *In vivo* experiments have investigated how the oviduct and endometrium as well as their secretions are modified by the metabolic status of the maternal environment (8, 9, 15, 17). While these reduced blood glucose and insulin concentrations alone are not detrimental for fertility and embryo survival (33) they result in a modified uterine environment that has consequences for offspring health (15, 34). Lactation results in a complex metabolic environment which not only has low concentrations of glucose and insulin but elevated concentrations of non-esterified fatty acids (NEFA), betahydroxybutyrate (BHB). Up to now it has not been possible to determine what role specific nutritional stressors play in modifying uterine function. Most *in vitro* models of bovine endometrium use traditional static cell culture or explants (26, 35–38) and while microfluidics have been used to recapitulate the human menstrual cycle (25) or the bovine oviduct (24), to our knowledge this is the first paper to report the use of microfluidics to mimic the bovine endometrium. This allowed us to determine the mechanism by which individual nutritional stressors alter endometrial function.

In early development, mammalian embryos use glucose as the main energy source to synthesize glycogen, nucleic acids, proteins, and lipids (39, 40). It is critical therefore that there is sufficient glucose transported into the uterine lumen to support embryo development however glucose can also act to modify the transcriptional response of a cell which modify the uterine environment. We demonstrated that exposure to altered concentrations of glucose changed the expression of transcripts involved in the biological processes of collagen fibril organization, blood vessel development, regulation of interferon α and β production, positive regulation of MAPK and ERK1/2 and positive regulation of energy homeostasis as well as the molecular functions of extracellular matrix (ECM) structural constituent, platelet-derived growth factor binging. Platelet activation is a complex signaling pathway positively dependent from several components as well as glycoprotein (GP) Ib-IX-V complex (GPIb-IX), phosphoinositide 3-kinase (PI3K-Akt), immunoreceptor tyrosine-based activation motif (ITAM), mitogen-activated protein kinase (MAPK), extracellular signal-regulated kinases 1 and 2 (ERK1/2), among others (41). Interactions between cells such as epithelial and stromal cells of the endometrium, involve mechanisms and extracellular matrix (ECM) constituents, including cell surface receptors (integrins) and receptors for fibronectin, collagen, and laminin. Stimulation of suspended platelets is an event dependent of collagen and thrombin and increase in intracellular Ca^2+^ is a key element in this process. Some of these events were connected with the GO terms involved with the DEGs when the endometrial cells were exposed to physiological extremes of glucose. Exposure to different concentrations of glucose not only altered DEGs in endometrial cells, they also altered the protein composition of *in vitro* ULF. This included proteins involved in Platelet Activation and Lysosome pathways.

Platelet activation plays a critical role in the function of platelets and it is involved in coagulation and inflammatory processes. Normally, platelet activation is induced by collagen or soluble platelet agonists that bind to G protein receptors, which stimulates the activation of platelet receptors (integrin α_IIb_β_3_), mediating platelet adhesion and aggregation (41). Interestingly, the Platelet-activating factor (PAF), a potent lipid mediator of inflammation and allergy, is involved in several reproductive processes (42) and PAF receptors are located in the oviduct of hamsters (43), mice (44) and oviduct and endometrium of cows (45). In humans, PFA increases vascular permeability and vasodilation, necessary processes for embryo implantation and plays an important paracrine role in stromal and epithelial cells interactions during this process (42). We have shown components of this pathway are altered by glucose and propose that may contribute to reduced endometrial function associated with altered glucose concentrations.

The other signaling pathway related with the proteins from the *in vitro* ULF is the Lysosome pathway. It is the primary site of cell digestion, lysosomes support cell function, recycling and providing a set of metabolites, such as amino acids, saccharides, lipids, ions and nucleobases and a key integrator and organizer of cellular adaptation and survival (46). Lysosomes link important metabolic processes encoded by AMPK (adenosine monophosphate-activated protein kinase) and GSK3β (glycogen synthase kinase 3) signaling hubs. AMPK is a primary cellular sensor for energy stress and glucose levels, promoting catabolic programs in response to low energy levels (47). AMPK has been linked to endometrial cancer in humans and depending on the context, AMPK can promote proliferation or cellular death (48). In addition, GSK3β is a kinase with apparently contradictory functions (49); for example, the presence of GSK3β in lysosomes can stimulate cell growth and survival, while its presence in the nucleus can promote cell death functions (50). In humans, GSK3β is expressed in endometrial cells and GSK-3β phosphorylated form presents cyclic variation, while phosphorylation of GSK-3β is regulated by progesterone and the inhibition of GSK-3β is temporally and related with the increased glycogen synthesis in the endometrial cells during the luteal phase (40, 51). Thus, the incorrect inactivation of GSK-3β could result in inadequate glycogen production and potentially to affect embryo implantation (40). In a previous study with dairy cows, increasing the circulating energy substrate, by exogenous infusion of glucose, was directly associated with a decrease in embryo development (size, width and area) and conception rates (34) contradictory to what was expected. We propose that the physiological glucose extremes may be directly affecting these important signaling pathways (Platelet activation and Lysosome) and interfering in processes such as receptivity and embryonic implantation.

Insulin has an essential metabolic role in regulating energy homeostasis in the body and insulin-dependent signaling also plays key functions in reproductive events and early development. In cattle, insulin concentrations vary throughout the estrous cycle (52), however, when an embryo is present in uterus, there is a decline insulin concentration in ULF (53). The bovine pre- and peri-implantation embryo expresses receptors for IGF and insulin receptors (54, 55) indicating the embryo is sensitive to concentrations of insulin during pre-implantation development (53). Depending on the nutritional status of the cow, insulin is suggested to have a metabolic function, regulating glucose levels into the uterus (56). Microfluidic exposure of endometrial cells to different concentrations of Insulin *in vitro* in this study altered signaling pathways in both epithelial and stromal cells related to metabolism. *In vivo*, circulating low levels of insulin are associated with negative energetic balance and the consequence for these low concentrations range from a delay in ovulation to an unfavourable environment for embryo development and even for the gestation maintenance (53). On the other hand, high levels of insulin although favourable for ovulation, are detrimental to the early embryo development (57, 58). Insulin supplementation during the *in vitro* development have had contradictory results with some improvement to morula stage embryos reported and some increase in cell number in the blastocysts, but most authors did not observe any effects on the blastocyst rate – a key developmental checkpoint (59–64). Even when rates of blastocyst development are not observed, an improvement in the number of cells in embryos could indicate a beneficial effect to the establishment of pregnancy. It occurs since blastocysts are able to produce and release embryotropins, involved in modulating endometrium transcripts mediated by interferon-tau and prostaglandin metabolism (65, 66). However, to the best of our knowledge, the insulin effects during the early luteal phase were not yet performed in *in vitro* bovine endometrial cells.

We observed that components of the complement and coagulation cascade were altered when our endometrium on-a-chip was exposed to low concentration of insulin. The complement system involved in the innate immune system and mostly functions to remove pathogens, dead cells, and debris. During early pregnancy there are changes to immune cells and markers in the endometrium (67, 68). In ruminants, receptivity to implantation involves a set of orchestrated events, among which the suppression of genes for immune recognition of the conceptus (embryo/fetus and associated membranes) and the increase in vascularization of the endometrium (Reviewed by (69)). Exposure to different concentration of insulin may modify these processes and contribute to dysregulated endometrial function.

Exposure of endometrial cells to physiological extremes of insulin also changed the composition of proteins in the *in vitro*-ULF. In general, the proteins up-regulated in low insulin concentration were associated with signaling pathways mainly related to metabolic events (Carbon metabolism, Biosynthesis of amino acids, Biosynthesis of antibiotics and Metabolic pathways). The proteins down-regulated when the endometrial cells were exposed to high concentrations of insulin were associated with Protein digestion and absorption, ECM-receptor interaction and Proteoglycans in cancer. Proteoglycans (including Syndecan-4 which we observed in our study) are often found on the cell surface or in the extracellular matrix (ECM) and perform multiple functions in cancer and angiogenesis by their ability to interact with both ligands and receptors that regulate neoplastic growth and neovascularization (70). Syndecan-4 connects two important signaling pathways (Proteoglycans in cancer and ECM-Receptors interactions) and it has been reported that blocking interactions between syndecan-4 and fibronectin decreases focal adhesions in cells, leading to increased cell proliferation in tumors. Thus, syndecan-4 has a key role in regulating cell adhesion, migration and proliferation in some tumors (71). The protein thrombospondin-1 (THBS1) was also associated with the ECM-receptor interaction pathway. This glycoprotein can bind to fibrinogen, fibronectin, laminin, collagen types V and VII and integrins mediating the interactions between cells and ECM. Also, THBS1 is involved in regulation of angiogenesis (72).

In humans, changes in ECM and/or in ECM-related signaling pathways are often attributed to pathological events, among which premature birth, cervical incompetence, endometriosis, polycystic ovary syndrome and neoplasms in the reproductive tract (73). Thus, demonstrating that ECM-receptor interactions pathway can be also involved in other tissues and biological events. In sheep and other livestock animals was previously reported, that the elongation of the conceptus requires a connection between the trophectoderm and the uterine luminal epithelium that causes a mosaic of interactions between integrins and extracellular matrix proteins (ECM), which act together to promote adhesion during implantation (74, 75). While our investigations were carried out in the pre-elongation stage of development, insulin-regulated changes to ECM components of the endometrium in early development may contribute to a compromised uterine environment and pregnancy loss.

In summary, we are the first to report the use of microfluidics to mimic the bovine endometrium *in vitro*. We demonstrate that this approach allows us to determine how individual components of maternal metabolism alter endometrial function. We specifically demonstrate at the transcriptional and proteomic levels that altered concentrations of glucose and insulin change ECM components of the endometrium. We propose that changes to these ECM components contribute to the compromised uterine environment associated with metabolic extremes in maternal circulation.

## ACKNOWLEDGMENTS

This work was supported by BBSRC grant number BB/R017522/1, QR-GCRF, and FAPESP (2016/22790-1 and 2018/14137-1). We would like to acknowledge the assistance of Stefania Mountevedi in helping with the bovine cell isolation and culture. We would like to thank Dr Ian Carr, Morag Raynor and Ummey Hany from the University of Leeds’s next generation sequencing facility core for undertaking sequencing analyses and the University of Bristol Proteomics core facility. We acknowledge Biorender in our production of components of Figure 1, and Pizza Loco & Rabbit Hole coffee for providing space, caffeine, and carbohydrates that helped fuel the writing of this manuscript during a global pandemic.

## Notes

### Competing Interest Statement

The authors have declared no competing interest.

